# Levetiracetam and Rufinamide are effective at supressing spike and wave seizure activity in an *in vitro* model of absence epilepsy

**DOI:** 10.1101/298711

**Authors:** Virdziniha Todorova, Georgina Ford, Roger D. Traub, Miles. A. Whittington, Stephen. P. Hall

**Affiliations:** Hull York Medical School, University of York, Heslington, YO10 5DD, UK; Dept. of Physical Sciences, IBM T.J. Watson Research Center, Yorktown Heights, NY 10598, USA.

**Keywords:** Absence epilepsy, Anti-epileptic Drugs, Spike-and-Wave, Levetiracetam, Rufinamide

## Abstract

Spike-and-wave discharges (SpW) are seen in absence-type epilepsies. They are heterogeneous in terms of their clinical burden and their electrographic signature, which is used to classify different types of absence seizures; typical absence, in which SpW frequency is 3-4Hz and atypical absence, which shows a slower 1-2Hz frequency. Treatment of SpW varies dependent upon the syndrome, but both Valproic Acid (VPA) and Ethosuximide (ESM) are shown to be effective in controlling typical absence seizures. Other anti-epileptic’s (AED’s), Levetiracetam (LEV) and Rufinamide (RUF), have shown promise in treating absence epilepsies and their associated syndromes. Here we examine the efficacy of these AED’s on an *in vitro* model of SpW.

Both LEV and RUF show an effective reduction in both the number of SpW events and the spike component amplitude; VPA shows no effect, whilst ESM enhances the spike amplitude. Phenytoin exacerbates the SpW activity, increasing both the number of SpW, amplitude of the SpW and the number of spikes within each event. These data suggest that both LEV and RUF could be effective in the treatment of absence-type epilepsies. They also suggest this model could be an effective tool to test other AED’s aimed at treating atypical absence syndromes.

## Introduction

Spike and wave discharges (SpW) are associated with a wide range of epilepsy pathologies, in particular absence type epilepsies. The concurrent involvement of the thalamus and neocortex is long established (Steriade et al., 1985; 1993; Crunelli & Leresche, 2002) and several models detailing the dysfunction of this interplay have been shown in which aberrant outputs from thalamic nuclei are emphasised (Destexhe et al., 1993; Destexhe, 1998). However, other studies have described a neocortical focus, showing that abnormal activity in the neocortex comes prior to that in the thalamus (Seneviratne et al., 2014). Our recent work (Hall et al, 2015) developed a model of SpW discharges in an isolated neocortical preparation, highlighting that neocortical networks alone are capable of generating SpW activity.

SpW events are known to be heterogeneous, which means that the SpW frequency, rhythmicity, amplitude distribution and clinical accompaniment can vary dramatically among different seizure pathologies (Blumenfeld, 2005). As such, absence epilepsies which present with SpW are split into two categories; typical absence and atypical absence. Typical SpW is usually described as being a 3-4Hz generalized SpW which lasts 10s of seconds (Gibbs et al., 1935) and are thought to arise from ‘normal’ sleep spindles (Kostoupolous, 2000). Atypical SpW have been shown at a slower frequency of around 1-2Hz and can be seen in secondary generalized epilepsies, such as Lennox-Gastaut syndrome (Niedermeyer, 1968). Whilst clinically there is a challenge to this delineation (Holmes et al., 1987), the roles of both thalamic and neocortical activity in typical and atypical absences is to be debated (Blumenfeld, 2005). Removal of the thalamus, cortex or thalamocortical tract in feline models of 3-4Hz generalized seizures leads to the ablation of SpW activity (Avoli & Gloor, 1981; 1982; 1982). In contrast, it has been shown that the injection of bicuculline to athalamic rats induces 2-4Hz SpW activity, whilst the injection of bicuculline direct to thalamus does not induce SpW seizures (Steriade & Contreras, 1998).

The treatment of SpW depends upon the underlying disease pathology but for typical 3-4Hz absence SpW, both Valproic acid (VPA) and Ethosuximide (ESM) are effective (Sato et al., 1982), as is Lamotrigine (Schlumberger et al., 1994). Both VPA and ESM have been shown to be more effective than Lamotrigine in controlling 3-4Hz typical absence seizures without intolerable side effects (Panayiotopoulos, 2001; Glauser et al., 2010) and as such are currently first treatment options for absence seizures in the NICE guidelines, with Lamotrigine to be considered when both VPA and/or ESM are ineffective or intolerable (NICE Clinical Guidelines, 2012). However, the treatment of slower, 1-2Hz atypical absence SpW in diseases such a Lennox-Gastaut is less well described, with NICE guidelines suggesting the use of VPA, for which there is little evidence for its efficacy (Ferrie & Patel, 2009). Novel anti-epileptic drugs (AED’s) such as Clobazam (Ng et al, 2011), Rufinamide (Kluger & Bauer, 2007; Resnik et al., 2011) and Levetiracetam (Verrotti et al., 2008; Fattore et al., 2011) have all shown efficacy in the treatment of childhood and juvenile absence epilepsy syndromes and as such Levetiracetam (LEV) and Rufinamide (RUF) are now listed in the NICE guidelines as therapies to be recommended through referral for absence seizures and for Lennox-Gastaut treatment (NICE Clinical Guidelines, 2012).

Considering the involvement of both the thalamus and the neocortex in clinical SpW, we wanted to examine the pharmacological profile of our neocortical isolated SpW model (Hall et al., 2015). By examining the efficacy of both established typical absence AED’s (VPA and ESM) and novel absence AED’s (LEV and RUF), combined with examining the effect of Phenytoin (PHE), an AED specifically described as do not prescribe in absence epilepsies (NICE Clinical Guidelines, 2012), we aimed to establish our *in vitro* SpW model as a specific absence type.

## Materials and Methods

### In Vitro electrophysiology

All surgical procedures were performed in accordance with the regulations of the United Kingdom Animals (Scientific Procedures) Act, 1986. Secondary somatosensory/parietal area coronal slices (450μm thick) were prepared from adult male Wistar rats (~200g) purchased from Charles River UK. All experiments were carried out between 10am and midday and rats were anaesthetised using isoflurane before terminal anaesthesia was conducted using a cocktail of xylazine and ketamine. A cardiac perfusion was performed before removal of the brain occurred. Slice were maintained at 34°C in a standard interface recording chamber containing oxygenated ACSF consisting of (in mM): 126 NaCl, 3 KCl, 1.25 NaH_2_PO_4_, 2 MgSO_4_, 1.2 CaCl_2_, 24 NaHCO_3_ and 10 glucose. Persistent, spontaneous delta oscillations were generated as described previously (Carracedo et al., 2013). Once the oscillations were deemed stable (3 recordings over 30 minutes in which the amplitude and frequency did not differ by more than 25%) further compounds could be bath applied.

Extracellular field recordings were obtained using micropipettes (2-5 MΩ) filled with ACSF and were bandpass filtered at 0.1Hz to 200Hz. Power spectra were derived from Fourier transform analysis of 60s epochs of data and results were presented as mean ± SEM. Spike detection and measurement was performed using Axograph and on the basis of transient deflections from the mean membrane voltage.

### Statistical analysis

was performed using SigmaPlot 12.3 (Systat Software Inc.). Normality of the data was tested using the Shapiro-Wilk normality test. For parametric data t-tests were used to measure any differences between the means of data before and after a manipulation. If the difference for more than one group of data was to be calculated a repeated measures Analysis of Variance (ANOVA) Test was used. For non-parametric data, a Mann-Whitney Rank Sum test was used. If the difference for more than one group of data was to be calculated a repeated measures Analysis of Variance (ANOVA) on Ranks was used.

### Materials

Perfusion of the following compounds was performed in certain experiments; (+)Tubocurarine chloride (10μM, antagonist to nicotinic acetylcholine and 5-HT_3_ receptors), BMS193885 (10μM, antagonist to NPY receptors), [D-p-Cl-Phe6, Leu17]-VIP (1μM, Agonist to VIP receptors), Levetiracetam (5-20μM, binds to SV2A), Rufinamide (5-50μM, Broad spectrum anti-convulsant), Phenytoin (5-20μM, blocks voltage gated sodium channels), Valproic Acid, sodium salt (5-20μM, Blocks voltage gated sodium channels), Ethosuximide (5-20μM, blocks T-type calcium channels). Drugs were obtained from Tocris Bioscience or Sigma Aldrich.

## Results

### The effect of AED’s upon delta rhythms in the secondary somatosensory cortex

The incidence of SpW has been shown to markedly increase with sleep (Angeleri et al., 1968). Previous work has highlighted that generating SpW *in vitro* is exquisitely dependent upon a background of delta frequency oscillations (Hall et al., 2015). Therefore, in order to examine the efficacy of AED’s on our model, it was first imperative to examine their effect upon delta oscillations.

The bath application of 5μM, 10μM or 20μM of ESM (Fig. 1A) showed no significant effects (P>0.05. n=5 slices, 3 animals for all concentrations) upon the amplitude (5μM; 142.45μV ± 82.36 vs. 189.3μV ± 106.65, 10μM; 195.96μV ± 76.74 vs. 172.48μV ± 69.58, 20μM; 402.28μV ± 131.98 vs. 499.48μV ± 163.09) or the frequency (5μM; 1.46Hz ± 0.1 vs. 1.46 Hz ± 0.09, 10μM; 1.32Hz ± 0.11 vs. 1.25 Hz ± 0.1, 20μM; 1.27Hz ± 0.3 vs. 1.27 Hz ± 0.1) of the delta oscillations. The bath application of 5μM, 10μM or 20μM of VPA (Fig. 1B) again showed no significant effects (P>0.05. n=5 slices, 3 animals for 5μM; 6 slices, 3 animals for 10μM and 20μM) upon the amplitude (5μM; 247.98μV ± 137.28 vs. 206.9μV ± 129.87, 10μM; 154.9μV ± 77.14 vs. 164.12μV ± 66.74, 20μM; 100.87μV ± 28.65 vs. 138.97μV ± 40.16) or the frequency (5μM; 1.41Hz ± 0.13 vs. 1.41 Hz ± 0.1, 10μM; 1.63Hz ± 0.15 vs. 1.63 Hz ± 0.07, 20μM; 1.54Hz ± 0.12 vs. 1.39 Hz ± 0.1) of the delta oscillations.

**Figure 1:**
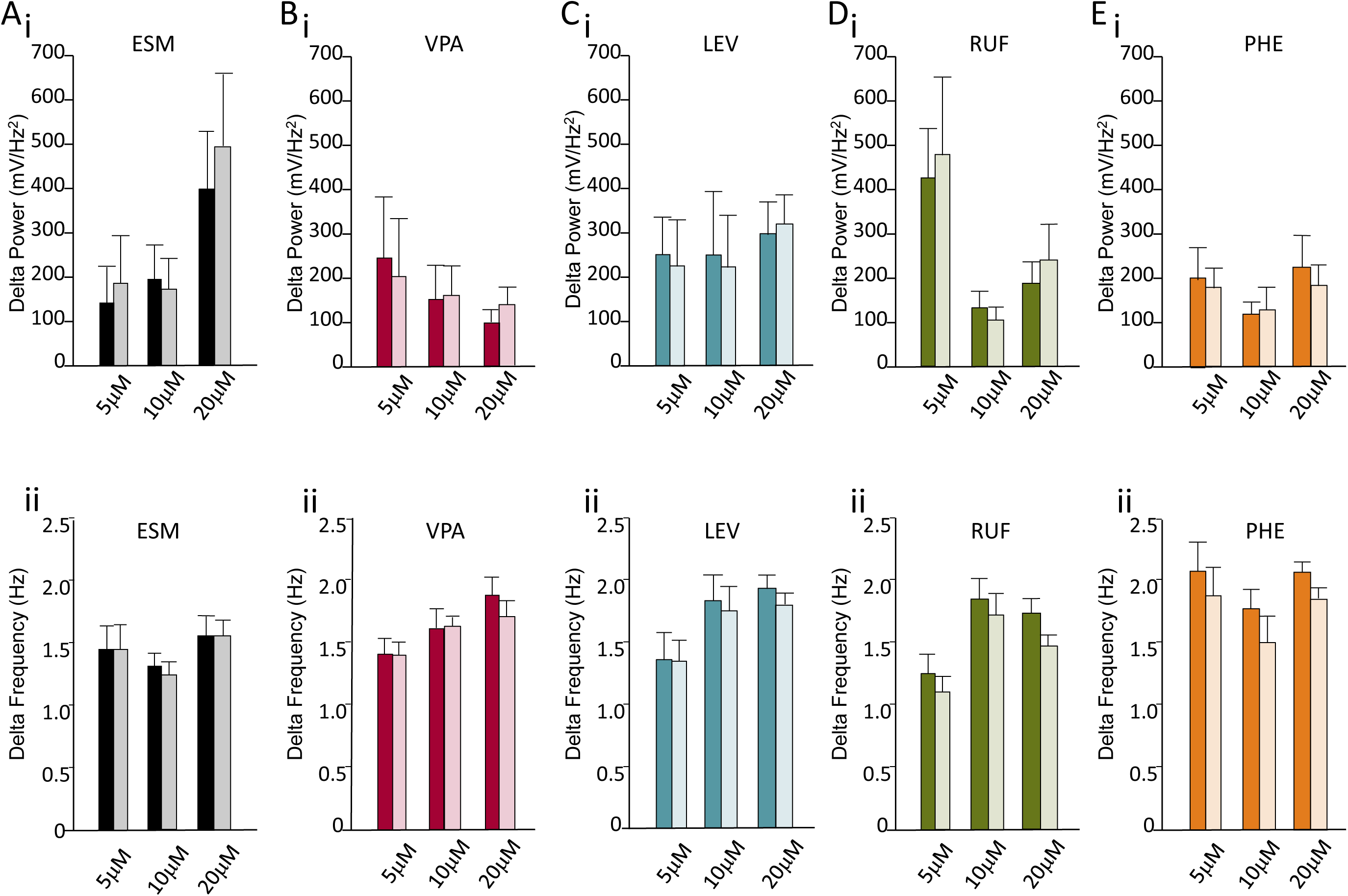
VPA, ESM, LEV, RUF and PHE have no effect on delta amplitude or frequency. **Ai.** Bar chart showing ESM had no effect upon delta amplitude at any of the concentration points. Solid bar = before ESM, transparent bar = after addition of ESM. **Aii.** Bar chart showing ESM had no effect upon delta frequency at any of the concentration points. Solid bar = before ESM, transparent bar = after addition of ESM. **Bi.** Bar chart showing VPA had no effect upon delta amplitude at any of the concentration points. Solid bar = before VPA, transparent bar = after addition of VPA. **Bii.** Bar chart showing VPA had no effect upon delta frequency at any of the concentration points. Solid bar = before VPA, transparent bar = after addition of VPA. **Ci.** Bar chart showing LEV had no effect upon delta amplitude at any of the concentration points. Solid bar = before LEV, transparent bar = after addition of LEV. **Cii.** Bar chart showing LEV had no effect upon delta frequency at any of the concentration points. Solid bar = before LEV, transparent bar = after addition of LEV. **Di.** Bar chart showing RUF had no effect upon delta amplitude at any of the concentration points. Solid bar = before RUF, transparent bar = after addition of LEV. **Dii.** Bar chart showing RUF had no effect upon delta frequency at any of the concentration points. Solid bar = before RUF, transparent bar = after addition of RUF. **Ei.** Bar chart showing PHE had no effect upon delta amplitude at any of the concentration points. Solid colour bar = before PHE, transparent bar = after addition of PHE. **Eii.** Bar chart showing PHE had no effect upon delta frequency at any of the concentration points. Solid bar = before PHE, transparent bar = after addition of PHE.

The bath application of 5μM, 10μM or 20μM of LEV (Fig. 1C) showed no significant effects (P>0.05. n=6 slices, 4 animals for 5μM; 7 slices, 5 animals for 10μM and 20μM) upon the amplitude (5μM; 254.0μV ± 83.28 vs. 228.62μV ± 105.58, 10μM; 252.3μV ± 143.26 vs. 226.99μV ± 118.04, 20μM; 303.27μV ± 76.15 vs. 322.18μV ± 66.59) or the frequency (5μM; 1.35Hz ± 0.24 vs. 1.34 Hz ± 0.17, 10μM; 1.82Hz ± 0.21 vs. 1.74 Hz ± 0.2, 20μM; 1.57Hz ± 0.09 vs. 1.46 Hz ± 0.07,) of the delta oscillations.

The bath application of 5μM, 10μM or 20μM of RUF (Fig. 1D) again showed no significant effects (P>0.05. n=6 slices, 4 animals for all concentrations) upon the amplitude (5μM; 427.61μV ± 110.17 vs. 482.51μV ± 176.58, 10μM; 133.69μV ± 37.27 vs. 109.59μV ± 29.35, 20μM; 192.67μV ± 49.56 vs. 245.04μV ± 80.29) or the frequency (5μM; 1.26Hz ± 0.15 vs. 1.11 Hz ± 0.13, 10μM; 1.85Hz ± 0.17 vs. 1.73 Hz ± 0.16, 20μM; 1.42Hz ± 0.09 vs. 1.2 Hz ± 0.17) of the delta oscillations.

Finally, the bath application of 5μM, 10μM or 20μM of phenytoin (Fig. 1E) showed no significant effects (P>0.05. n=6 slices, 3 animals for all concentrations) upon the amplitude (5μM; 199.92μV ± 72.3 vs. 181.4μV ± 45.1, 10μM; 120.65μV ± 27.66 vs. 129.87μV ± 51.52, 20μM; 229.32μV ± 72.16 vs. 186.28μV ± 48.61) or the frequency (5μM; 2.08Hz ± 0.23 vs. 1.89 Hz ± 0.23, 10μM; 1.78Hz ± 0.17 vs. 1.51 Hz ± 0.21, 20μM; 1.69Hz ± 0.07 vs. 1.52 Hz ± 0.15) of the delta oscillations.

### The effect of established AED’s on SpW activity in the secondary somatosensory cortex

The NICE guidelines suggest the use of ESM, VPA and lamotrigine, either in mono-therapy or in combination therapy to supress absence type seizure. Studies have highlighted the greater efficacy and reduced side effects offered by ESM and VPA when compared to lamotrigine (Glauser et al., 2010; 2013). Given this, we decided to examine the effect of ESM and VPA upon SpW activity, generated as per Hall et al., 2015. On a background of persistent delta oscillations we generated SpW activity in both control conditions, with the bath application of d-TC alone and by bath applying d-TC in the presence of varying concentrations of ESM and VPA. We then examined the number of SpW events per minute and the amplitude of the spike component of the SpW event.

Bath application of 5μM, 10μM or 20μM ESM prior to the generation of SpW activity failed to supress seizure activity (Fig. 2 A, B, Ei). At 5μM, the number of SpW events per minute did not significantly differ, averaging 10.29min^−1^ ± 0.67 with d-TC alone and 11.00min^−1^ ± 1 with ESM (P>0.05, n=5 slices/3 animals). Again, no difference was observed at either 10μM ESM (11.40min^−1^ ± 0.92; P>0.05, n=5 slices/3 animals) or 20μM ESM (11.60min^−1^ ± 0.36; P>0.05, n=5 slices/3 animals). The amplitude of the spike component did show a significant increase at 5μM ESM, averaging at 423.29μV ± 46.96 with d-TC alone and 769.78μV ± 92.4 with ESM (P=0.016, n=5 slices/3 animals). This increase was sustained but non-significant at 10μM (608.2μV ± 184.29; P>0.05, n=5 slices/3 animals), but was significant once more at 20μM (740.6μV ± 83.13; P=0.022, n=5 slices/3 animals).

**Figure 2:**
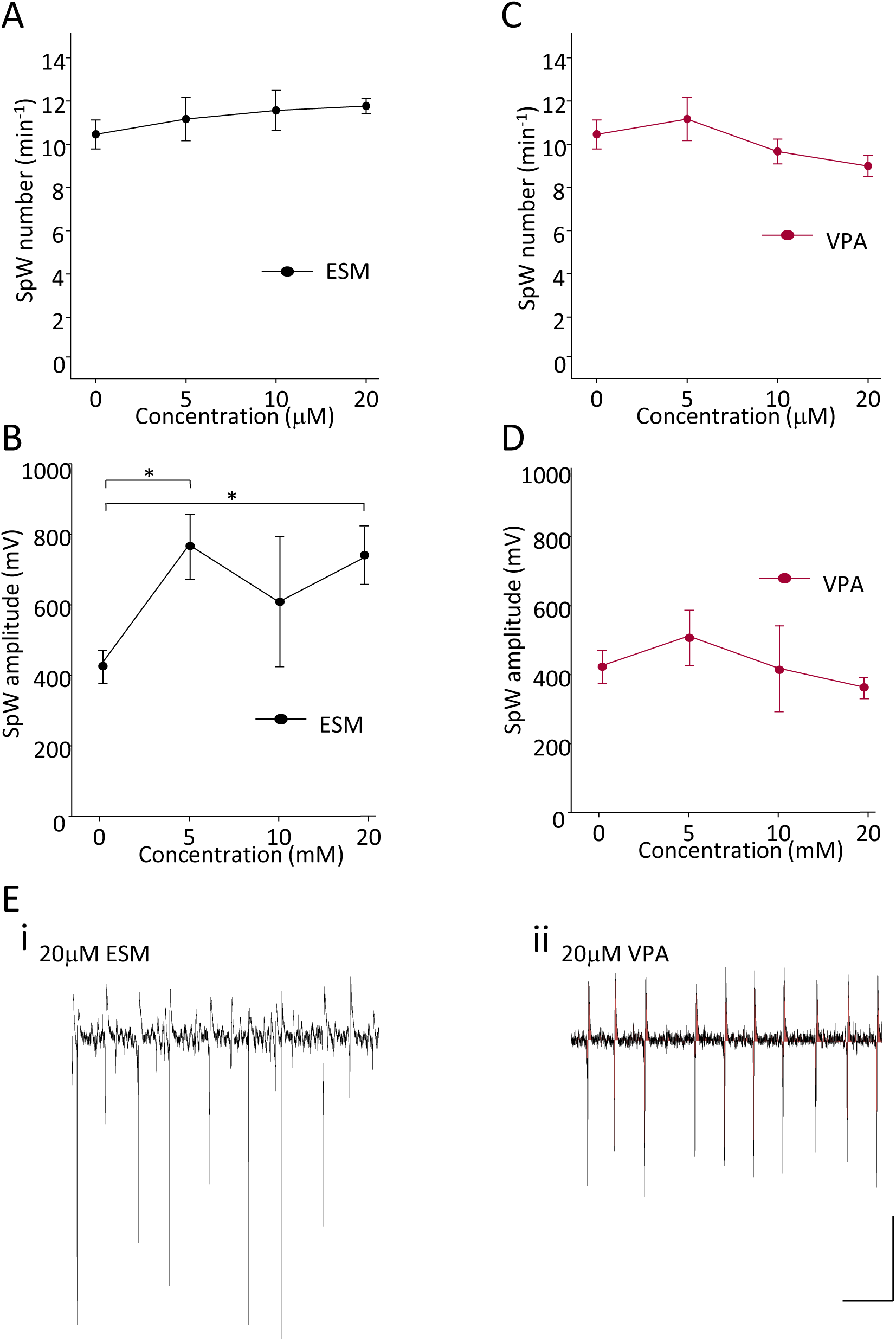
ESM and VPA show no suppression of d-TC induced SpW activity. **A.** Line graph showing ESM has no effect upon SpW number. **B.** Line graph showing ESM has an exacerbating effect upon the amplitude of the spike component of SpW at 5 and 20μM. **C.** Line graph showing VPA has no effect upon SpW number. **D.** Line graph showing VPA has no effect upon the amplitude of the spike component of SpW. **E.** Example trace showing SpW activity recorded from S2 in the presence of 20μM ESM. **F.** Example trace showing SpW activity recorded from S2 in the presence of 20μM VPA. Scale bar = 200μV, 10 seconds. * indicates P<0.05.

Bath application of 5μM, 10μM or 20μM VPA prior to the generation of SpW activity failed to supress seizure activity (Fig. 2 C, D, Eii). At 5μM, the number of SpW events per minute showed no significant change, averaging 10.29min^−1^ ± 0.67 with d-TC alone and 11.00min^−1^ ± 1.1 with VPA (P>0.05, n=5 slices/3 animals). Again, no difference was observed at either 10μM VPA (9.5min^−1^ ± 0.57; P>0.05, n=6 slices/3 animals) or 20μM VPA (8.83min^−1^ ± 0.48; P>0.05, n=6 slices/3 animals). The amplitude of the spike component also showed no significant change at 5μM ESM, averaging at 423.29μV ± 46.96 with d-TC alone and 512.23μV ± 78.78 with VPA (P>0.05, n=5 slices/3 animals). Again, no differences were observed at 10μM (419.08μV ± 125.17; P>0.05, n=6 slices/3 animals) or 20μM (361.17μV ± 29.6; P>0.05, n=6 slices/3 animals).

### The effect of novel AED’s on SpW activity in the secondary somatosensory cortex

LEV has been shown to provide seizure suppression and an increase in cognitive function in patients with Lennox-Gastaut syndrome (De Los Reyes et al., 2004; Diaz-Negrillo et al., 2011) as has RUF (Resnik et al., 2011). Given this, we decided to challenge our SpW activity, generated as per Hall et al., 2015, with LEV and RUF. On a background of persistent delta oscillations we generated SpW activity in both control conditions, with the bath application of d-TC alone, and by bath applying d-TC in the presence of varying concentrations of LEV and RUF. We then examined the number of SpW events per minute and the amplitude of the spike component of the SpW event.

Bath application of all concentrations of LEV prior to the generation of SpW activity successfully supressed seizure activity (Fig. 3 A, B, Ei), averaging 10.29min^−1^ ± 0.67 with d-TC alone and 6.00min^−1^ ± 0.97 with LEV (P<0.05, n=6 slices/4 animals). At 10μM, LEV further reduced SpW activity (1.29min^−1^ ± 0.61; P<0.05, n=7 slices/5 animals) and at 20μM, LEV had completely supressed SpW events (0min^-1^ ± 0; P<0.05, n=5 slices/4 animals). The amplitude of the spike component did not show a significant change at 5μM LEV, averaging at 423.29μV ± 46.96 with d-TC alone and 761.11μV ± 301.62 with LEV (P>0.05, n=6 slices/4 animals). However, at 10μM there was a significant reduction in the amplitude of the spike (183.68μV ± 118.84; P<0.05, n=7 slices/5 animals) and given the SpW suppression, the lack of SpW meant a significant reduction at 20μM (0μV ± 0; P<0.001, n=7 slices/4 animals).

**Figure 3:**
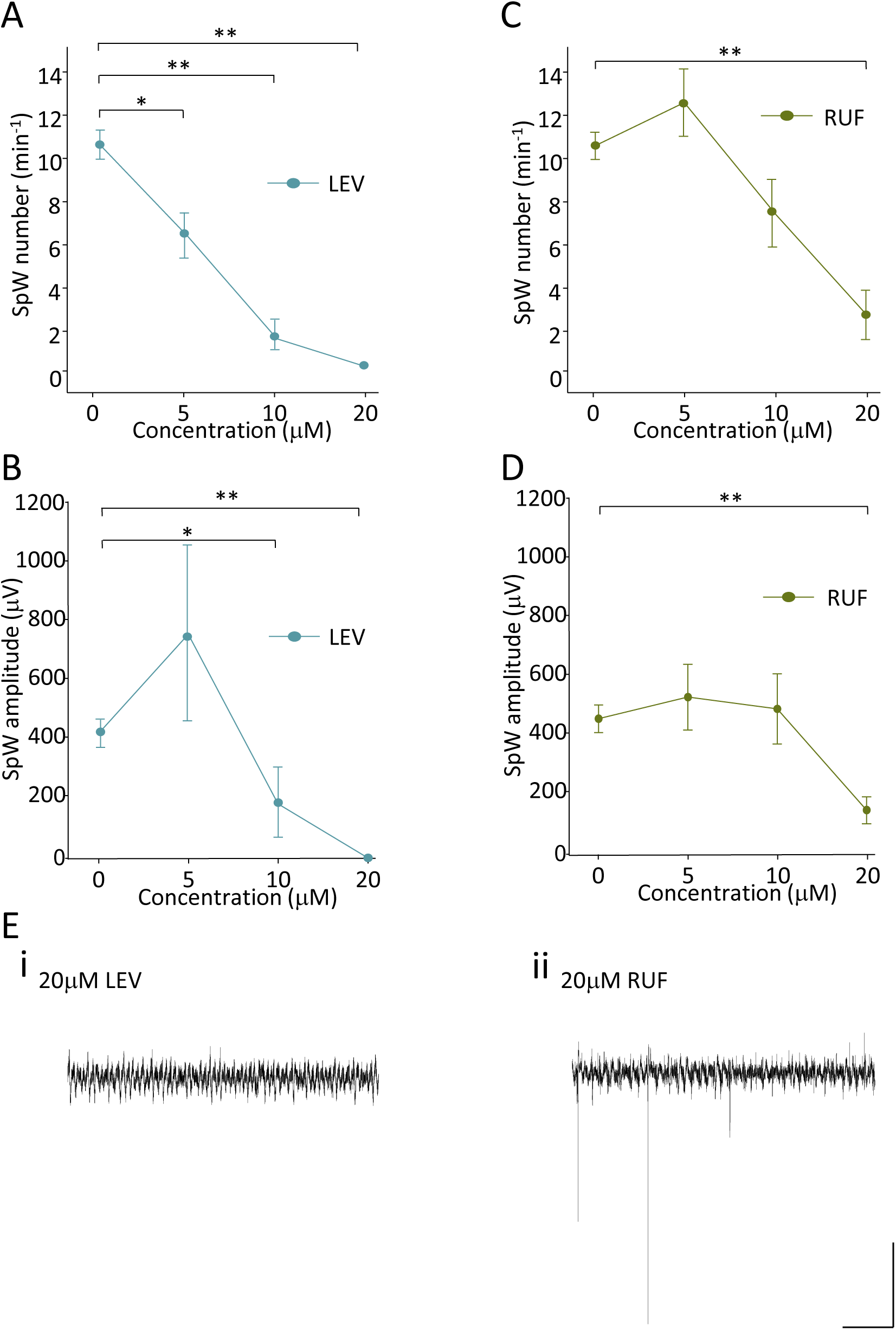
LEV and RUF show effective suppression of d-TC induced SpW activity. A. Line graph showing the reduction in SpW number, which correlates with increasing concentrations of LEV. B. Line graph showing LEV reduces the amplitude of the spike component of SpW at 5 and 20μM. C. Line graph showing RUF reduces SpW number at 20μM D. Line graph showing RUF shows a significant reduction in SpW amplitude at 20μM. E. Example trace showing SpW activity recorded from S2 in the presence of 20μM LEV. F. Example trace showing SpW activity recorded from S2 in the presence of 20μM RUF. Scale bar = 100μV, 10 seconds. * indicates P<0.05. ** indicates P<0.001.

Bath application of high concentrations of RUF prior to the generation of SpW activity also supressed seizure activity (Fig. 3 C, D, Eii). At 5μM, the number of SpW events per minute showed no significant change, averaging 10.29min^−1^ ± 0.67 with d-TC alone and 12.33min^−1^ ± 1.61 with RUF (P>0.05, n=6 slices/4 animals). Again, no difference was observed at 10μM RUF (7.17min^−1^ ± 1.61; P>0.05, n=6 slices/3 animals). However, at 20μM RUF there was significant suppression of SpW activity (2.5min^−1^ ± 1.15; P<0.05, n=6 slices/5 animals). The amplitude of the spike component showed no significant change at 5μM RUF, averaging at 423.29μV ± 46.96 with d-TC alone and 496.89μV ± 111.49 with RUF (P>0.05, n=6 slices/4 animals). Again, no differences were observed at 10μM (456.87μV ± 119.38; P>0.05, n=6 slices/3 animals), but at 20μM RUF there was a significant reduction in the amplitude of the spike (111.78μV ± 45.92; P<0.05, n=6 slices/5 animals).

### The effect of phenytoin on SpW activity in the secondary somatosensory cortex

PHE is currently listed as do not prescribe for absence type seizures (NICE guidelines 2012). We wanted to examine its effect upon our *in vitro* model of SpW events to determine if it had an exacerbating effect upon the SpW activity. On a background of persistent delta oscillations we generated SpW activity in both control conditions, with the bath application of d-TC alone and by bath applying d-TC in the presence of varying concentrations of PHE.

Bath application of high concentrations of PHE prior to the generation of SpW activity caused an increase to both SpW number and spike amplitude (Fig. 4 A, B, D). At 5μM, the number of SpW events per minute showed no significant change, averaging 10.29min^−1^ ± 0.67 with d-TC alone and 11.2min^−1^ ± 1.4 with PHE (P>0.05, n=5 slices/3 animals). Again, no difference was observed at 10μM PHE (9.33min^−1^ ± 1.31; P>0.05, n=6 slices/3 animals). However, at 20μM PHE there was a significant increase in SpW activity (15.5min^−1^ ± 2; P<0.05, n=6 slices/3 animals). The amplitude of the spike component showed significant change at 5μM PHE, averaging at 423.29μV ± 46.96 with d-TC alone and 797μV ± 62.6 with PHE (P<0.05, n=5 slices/3 animals). This increase in amplitude was enhanced further at 10μM PHE (1819.33μV ± 520.74; P<0.001, n=6 slices/3 animals), and at 20μM (1543.33μV ± 229.18; P<0.001, n=6 slices/3 animals).

**Figure 4:**
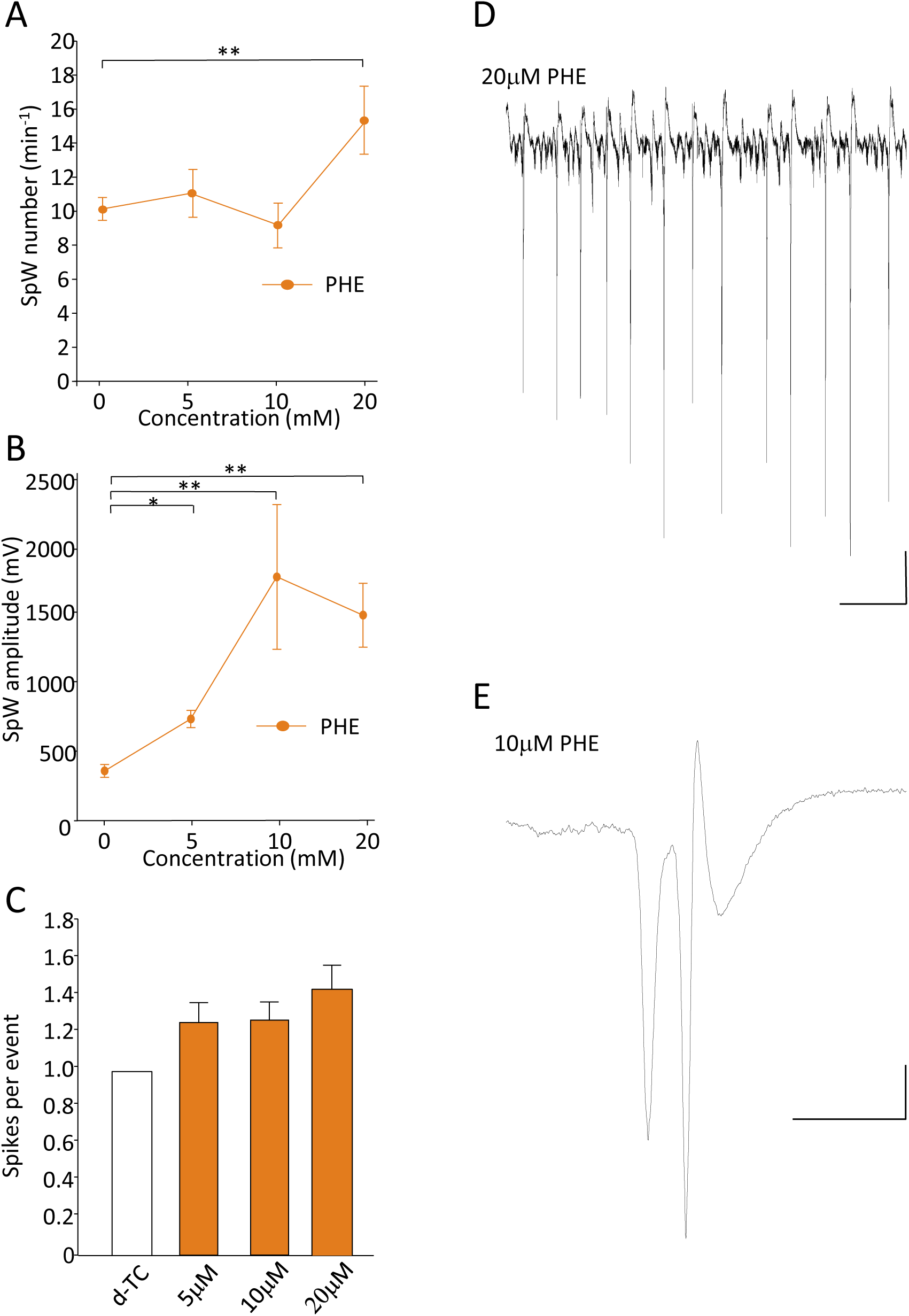
PHE exacerbates d-TC induced SpW activity. A. Line graph showing an increase in SpW number at 20μM PHE. B. Line graph showing PHE significantly increases the amplitude of the spike component of SpW at all concentration points. C. Bar chart showing an increase in the number of spikes per event when SpW were induced by d-TC in the presence of PHE. Clear bar show number of spikes per SpW event in d-TC alone, coloured bars in the presence of PHE. D. Example trace showing SpW activity recorded from S2 in the presence of 20μM PHE. Scale bar = 200μV, 10 seconds. F. Example trace showing a single SpW event recorded from S2 in the presence of 20μM PHE. Scale bar= 100μV, 200 milliseconds. * indicates P<0.05. ** indicates P<0.001.

In testing all other AED’s only a single spike was observed within each SpW event. However, with the bath application of PHE, instances of multiple spikes were observed within each SpW (Fig. 4 C, E). At 5μM PHE, the number of spikes per SpW event increased from 1 ± 0 with d-TC alone, to 1.27 ± 0.11 with PHE. This increase was maintained at 10μM PHE (1.28 ± 0.09) and at 20μM (1.45 ± 0.13). However, none of the increases in spike number were significant (P>0.05. n=5 slices/3animals for 5μM and 10μM, n=6 slices/3animals for 20μM).

### The effect of Levetiracetam on SpW activity induced by peptidergic disinhibition, in the secondary somatosensory cortex

In Hall et al, 2015, we explored the mechanisms behind the development of SpW. It was observed that SpW was caused by the disruption in the balance between two peptides, neuropeptide Y (NPY) and vasointestinal peptide (VIP). Using a combination of an NPY antagonist (10μM BMS193885) and 1μM VIP on a background of persistent delta oscillations, SpW was observed. The most overt effect of the AED’s on d-TC induced SpW activity was with LEV. We wanted to test the effect of LEV on the peptidergic model of SpW activity, to examine if its effect was maintained in this more physiological model.

Bath application of 20μM LEV prior to the generation of SpW activity caused a significant reduction in both SpW number (Fig. 5 A, B, D). The number of SpW events reduced from 16.29min^−1^ ± 1.19 with BMS 193855 and VIP and 1min^−1^ ± 0.43 with 20μM LEV (P<0.05, n=8 slices/4 animals BMS 193855/VIP, n=7 slices/4 animals 20μM LEV). Whilst the number of SpW per minute was reduced with the addition of LEV, the SpW amplitude was not significantly reduced. The amplitude of the spike component also showed a non-significant decrease from 106.8μV ± 20.92 with BMS 193885 and VIP and 70.75μV ± 20.13 with 20μM LEV(P>0.05, n=8 slices/4 animals BMS 193855/VIP, n=7 slices/4 animals 20μM LEV).

**Figure 5:**
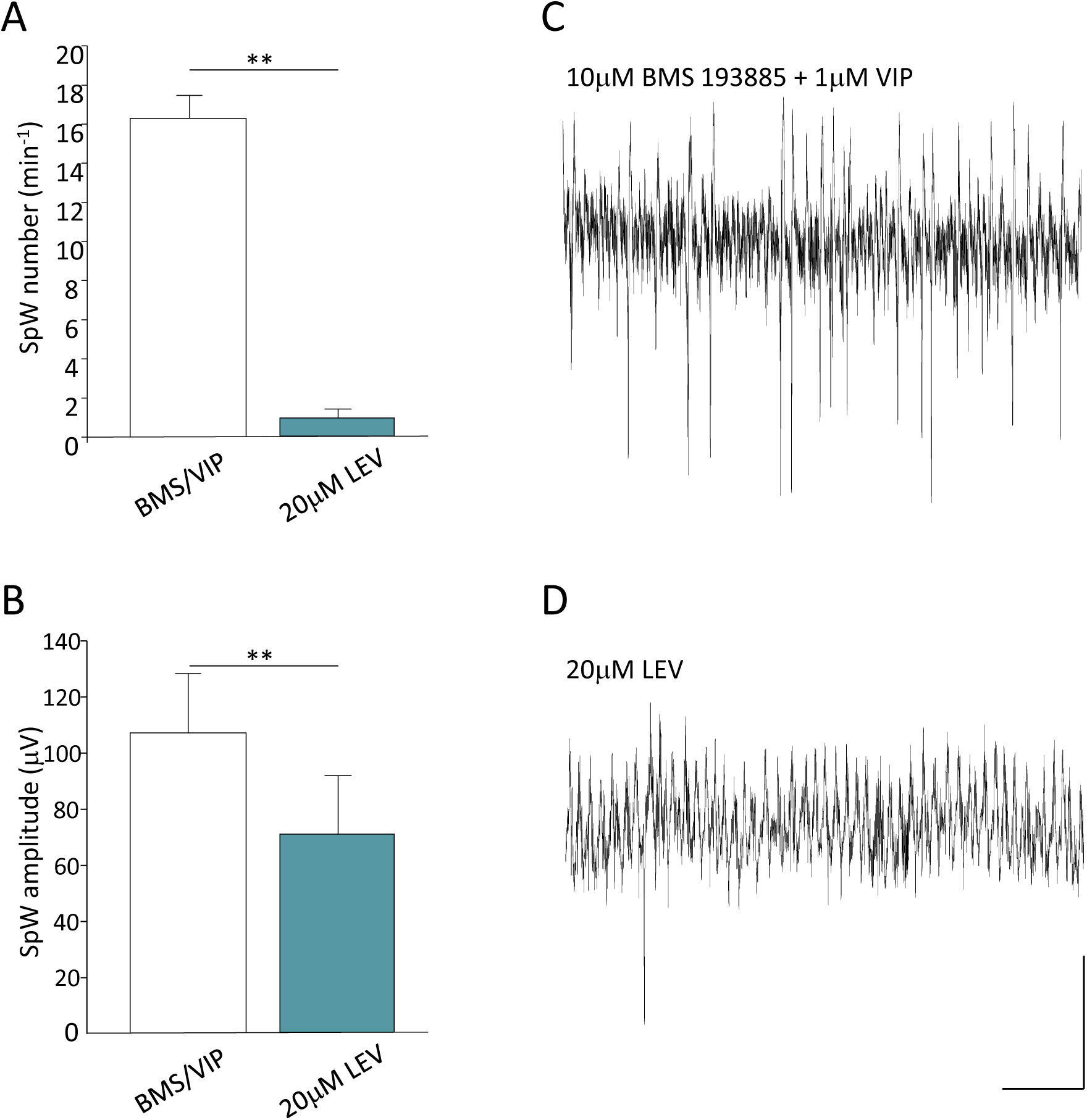
LEV supresses SpW induced by peptidergic disinhibition. A. Bar chart showing the reduction of SpW number by 20μM LEV, when SpW is induced by the peptidergic disinhibition model. Clear bar shows peptidergic disinhibition alone, blue bar show peptidergic disinhibition in the presence of 20μM LEV. B. Bar chart showing the reduction of SpW amplitude by 20μM LEV, when SpW is induced by the peptidergic disinhibition model. Clear bar shows peptidergic disinhibition alone, blue bar show peptidergic disinhibition in the presence of 20μM LEV. C. Example trace showing SpW activity recorded from S2 following peptidergic disinhibition. D. Example trace showing a SpW activity recorded from S2 following peptidergic disinhibition in the presence of 20μM LEV. Scale bar = 100μV, 10 seconds. PHE. Scale bar = 100μV, 200 milliseconds. * indicates P<0.05. ** indicates P<0.001.

## Discussion

The interplay between the thalamus and cortex in the development of SpW activity is long established. However, the role of the thalamus and/or cortex in different SpW types is complex. The removal of the thalamus, cortex or thalamocortical tract in feline models of 3-4Hz generalized seizures leads to the ablation of SpW activity (Avoli & Gloor, 1981; 1982; 1982), whilst in genetic absence epilepsy rats of Strasbourg (GAERS), faster SpW (c7-11Hz) cannot be sustained with the removal of either thalamus or neocortex (Danober et al., 1998). Furthermore, mean field modelling of the corticothalamic system shows that increasing the feed-forward inhibition of cortex to thalamic relay neurons can supress 3-4Hz SpW activity (Chen et al., 2017). These studies seem to suggest a role for both thalamus and neocortex in typical absence seizures. Conversely, SpW activity when the thalamus is ablated and when cortex is isolated has also been described (Steriade and Contreras, 1998; Timofeev & Steriade, 2004). However, some of the SpW activity within these models appears to be at the slower 1-2Hz frequency more associated with atypical absence. Given this evidence, it has been postulated that the involvement of the thalamus in the generation of SpW may depend upon the model (Blumenfeld, 2005). The model studied here is an isolated neocortical model; the thalamus plays no role in the generation of SpW.

SpW have been shown to increase with sleep (Angeleri et al., 1968) and the dependence upon delta frequency activity to generate SpW in this model was highlighted in Hall et al., 2015. The data within this study show that no AED’s tested have any significant effect upon the delta rhythm in vitro, allowing us to be sure we are testing their efficacy upon the SpW events. Both of the AED’s used to treat typical absence seizures, VPA and ESM, had no positive effect upon SpW activity. ESM, whilst not altering the number of SpW events observed, did increase the amplitude of the spike component of the events. The spike component of the event in this model has been shown to be generated from the superficial layers of the cortex (Hall et al., 2015). As such an increase in the amplitude of the spike component suggests a shift from cells firing single spikes, to firing bursts. Thus the increase in the spike amplitude suggests that ESM is exacerbating the SpW, akin to the effect of PHE in this model. PHE increased the SpW number at high doses and increased the amplitude of the spike component throughout the concentration range. More significantly PHE caused an increase in the number of spikes within each event at all doses, highlighting a distinct exacerbation of SpW as previously considered (Genton, 2000). As such it appears correct that PHE is on the do not prescribe list for absence type epilepsies.

LEV showed effective SpW reduction, both in term of SpW number and in terms of the spike component amplitude. Patient studies have shown LEV improves atypical absence seizures observed in disorders such as Lennox-Gastaut (De Los Reyes et al., 2004), Angelman syndrome (Shaaya et al., 2016) and Jeavons syndrome (Striano et al., 2008). It also has been shown to improve the cognitive deficits associated with Lennox-Gastaut syndrome (Diaz-Negrillo et al., 2011). However, it has been described as aggravating typical absence seizures (Auvin et al., 2011). The other AED to show effective reductions in SpW within this study was RUF. Again, RUF is effective in the treatment atypical absence seizures (Albini et al., 2016) particularly in diseases such as Lennox-Gastaut (Glauser et al., 2008) and as such it has recently been approved as an adjunctive therapy.

The model used in this study uses isolated neocortex to generate SpW and given the evidence discussed above the model could be described as being atypical in its presentation. A dramatic reduction in SpW observed with the application of LEV and RUF, which have shown efficacy in patients with atypical absence epilepsy but not in patients with typical absence, was observed.

Couple this with the lack of efficacy of VPA and ESM, which are both prescribed for typical absence epilepsies and the evidence would suggest that the *in vitro* model used in this study and first described in Hall et al, 2015 is of atypical absence epilepsy. Given the slower frequency of atypical absence epilepsies discussed in the introduction, this would potentially explain the slower frequency SpW activity observed in this model (Hall et al., 2015). The implications of this study are two-fold: 1) It further highlights the excellent efficacy of LEV and RUF in the treatment of atypical absence epilepsies; 2) Given the difficulties in treating atypical absence epilepsies (Sinclair and Unwala, 2007) this model could provide an effective tool to test novel AED’s aimed at treating atypical absence epilepsies.

## Acknowledgements

We thank The Wellcome Trust for funding this work in the UK. RDT was funded by IBM & NIH/NINDS R01NS044133.

